# An integrated comparative physiology and molecular approach pinpoints mediators of breath-hold capacity in dolphins

**DOI:** 10.1101/2021.01.20.425775

**Authors:** Ashley M. Blawas, Kathryn E. Ware, Emma Schmaltz, Larry Zheng, Jake Spruance, Austin S. Allen, Nicole West, Nicolas Devos, David L. Corcoran, Douglas P. Nowacek, William C. Eward, Andreas Fahlman, Jason A. Somarelli

## Abstract

Ischemic events, such as ischemic heart disease and ischemic stroke, are the number one cause of death globally. Ischemia prevents blood, carrying essential nutrients and oxygen, from reaching tissues and organ systems, leading to cell and tissue death, and eventual organ failure. While humans are relatively intolerant to these ischemic events, other species, such as marine mammals, have evolved remarkable tolerance to chronic ischemia/reperfusion during diving. Here we capitalized on the unique adaptations of bottlenose dolphins (*Tursiops truncatus*) as a comparative model of ischemic stress and hypoxia tolerance to identify molecular features associated with breath-holding. Using RNA-Seq we observed time-dependent upregulation of the arachidonate 5-lipoxygenase (ALOX5) gene during breath-holding. Consistent with the RNA-Seq data, we also observed increased ALOX5 enzymatic activity in the serum of dolphins undergoing breath holds. ALOX5 has previously been shown to be activated during hypoxia in rodent models, and its metabolites, leukotrienes, induce vasoconstriction. The upregulation of ALOX5 occurred within the estimated aerobic dive limit of the species, suggesting that ALOX5 enzymatic activity may promote tolerance to ischemic stress through sustained vasoconstriction in dolphins during diving. These observations pinpoint a potential molecular mechanism by which dolphins, and perhaps other marine mammals, have adapted to the prolonged breath-holds associated with diving.

## Introduction

### Ischemic stress and hypoxia are associated with negative clinical outcomes in humans

Maintenance of homeostatic function in mammalian tissues is directly dependent on a continuous supply of oxygenated blood. Interruption of this blood supply, known as ischemia, results in reduction in local oxygenation compared to normal physiologic levels, or hypoxia, and can lead to inflammation and cell/tissue death (Bona et al., 1999; Choi, 1996; Eltzschig and Carmeliet, 2011; Gottlieb and Engler, 1999; Murdoch et al., 2005). Ischemia is the causative factor in multiple clinical settings, and ischemic heart disease is the number one cause of death globally, accounting for over 9 million deaths each year (Nowbar et al., 2019; World Health Organization, 2018).

### Marine mammals have evolved tolerance to ischemic stress

While humans have little tolerance for ischemic stress and hypoxia, a number of species have evolved unique physiologies that allow them to seemingly thrive despite regular tissue-level ischemia and low-oxygen environments. One group of animals that undergo repeated daily ischemic events is marine mammals. During a dive, a marine mammal experiences a suite of cardiovascular changes that aid in reducing aerobic metabolism (Irving et al., 1941; Scholander, 1940). As part of this response, both heart rate (*f*_H_) and stroke volume decrease, resulting in reduced cardiac output (Fahlman et al., 2020b, 2019b). Increased peripheral resistance, through selective vasoconstriction, helps assure that mean arterial blood pressure is maintained, at least in studies on forced diving in seals (Blix et al., 1976; Zapol et al., 1979). Ultimately, this response conserves oxygen in the blood and lungs for oxygen-sensitive tissues like the brain and the heart, while the skeletal muscles rely on endogenous myoglobin-bound oxygen for aerobic metabolism (Davis and Kanatous, 1999; Fahlman et al., 2009). While these responses to submersion in water are largely conserved across all vertebrates, many of the physiological adaptations that support diving are exaggerated in marine mammals compared to other taxa (Kooyman and Ponganis, 1998; Panneton, 2013) (**Figure 1**). For example, maintenance of increased peripheral resistance does not appear to occur in human breath-hold divers, as mean arterial blood pressure increases with dive duration (Breskovic et al., 2011; Gooden, 1994; Taboni et al., 2019). These physiological differences highlight the tremendous potential to study marine mammals as model organisms for the investigation of adaptations to ischemic and hypoxic stress tolerance, and the cardiorespiratory plasticity that helps prevent hypertension (Blawas et al., 2021; Fahlman et al., 2019b, 2020b).

**Figure 1.**
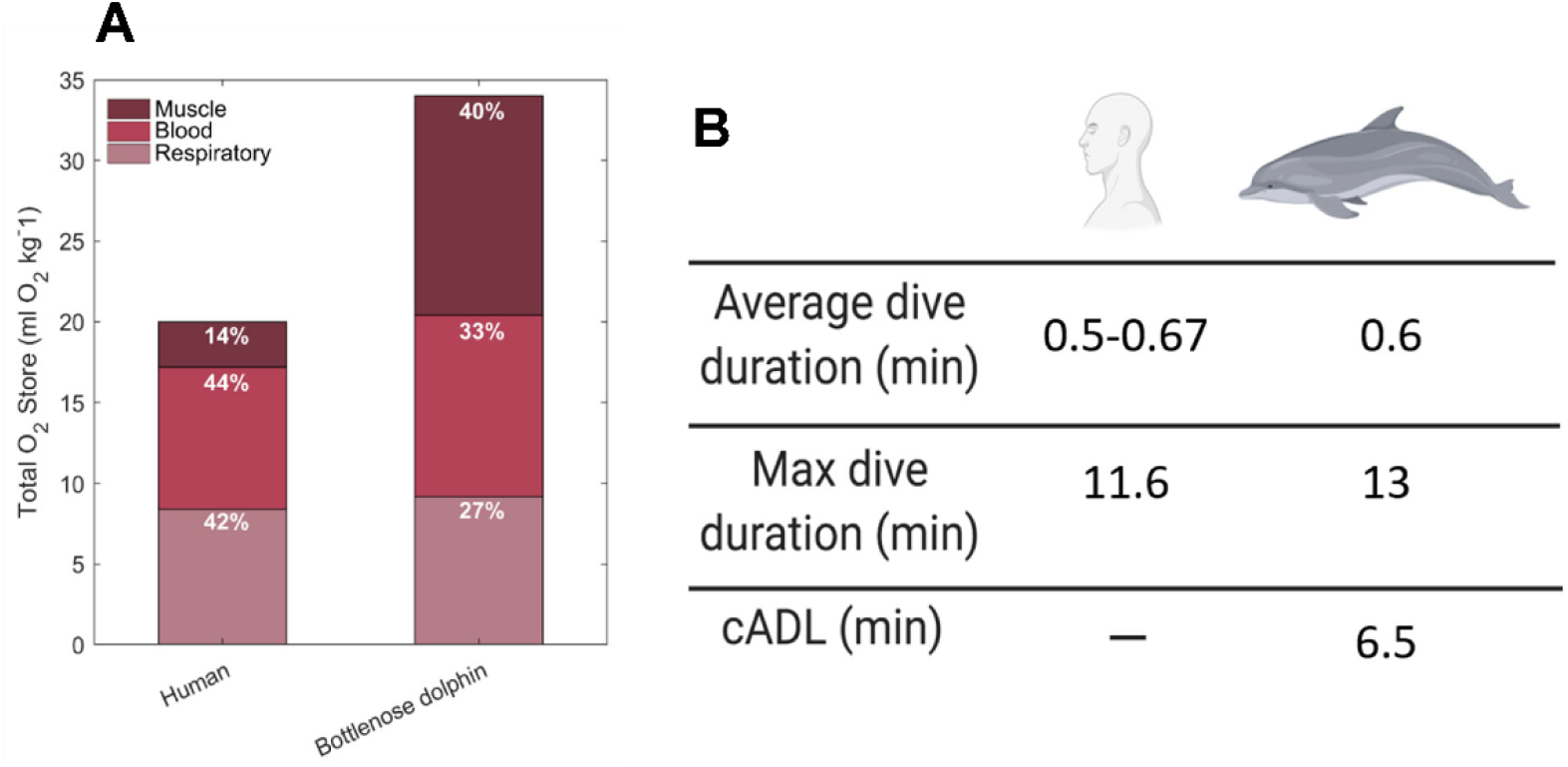
Dolphins as a model of ischemia. A. Dolphins and other cetaceans have increased oxygen stores that are reapportioned compared to humans. Oxygen store data were reported in Ponganis et al., 2011 (human) and Kooyman and Ponganis, 2018 (dolphin). B. The enhanced oxygen stores and diving capacity of dolphins makes them a unique model to study ischemic stress tolerance. Dive data and calculated aerobic dive limit (cADL) were reported by AIDA and Foster and Sheel, 2005 (human) and Fahlman et al., 2018 (dolphin).

### Marine mammals have evolved molecular adaptations to ischemic stress tolerance

Increasing attention has been paid to the defenses marine mammals possess against the oxidant by-products and inflammation associated with ischemic, hypoxia, and reperfusion at the molecular level (Allen and Vázquez-Medina, 2019; Hindle, 2020; Zhu et al., 2018). Using phylogenetic and evolutionary convergence approaches, several gene families have been identified to contribute to the increased ischemic stress tolerance of marine mammals, including hypoxia-inducible factor 1 (HIF-1) (Bi et al., 2015; Johnson et al., 2005, 2004), genes relating to the glutathione system and peroxiredoxins (Bagchi et al., 2018; Tift et al., 2014; Yim et al., 2014; Zhou et al., 2018), and several genes linked to oxygen storage, particularly hemoglobin and myoglobin (Mirceta et al., 2013; Nery et al., 2013; Tian et al., 2017, 2016). Yet, few studies have examined differential gene expression in marine mammals under conditions of ischemia and hypoxia (i.e. diving conditions).

Here, we investigate the dynamic molecular changes that occur during an apnea in bottlenose dolphins using genomic analysis of peripheral blood mononuclear cells (PBMCs) and serum sampled at regular intervals during breath-holds. We couple these analyses with previously-published *f*_H_ measurements from the same dolphins to understand how the timing of molecular changes relates to the physiologic dive response (Blawas et al., 2021; Fahlman et al., 2019b, 2020b). Our integrated analyses pinpoint a gene regulatory network centered around the arachidonate 5-lipoxygenase (ALOX5) gene and its downstream metabolites, leukotrienes, as differentially activated during breath-holding. This activation of ALOX5 is consistent with cardiovascular control through reduction in *f*_H_ and peripheral vasoconstriction to efficiently manage O2 use during diving. Based on our collective results we propose a model in which the ALOX5 pathway is upregulated during extended breath-holds as a mechanism to sustain vasoconstriction and maintain oxygen stores for critical organs while diving.

## Materials and Methods

### Data collection and animal information

Four adult male bottlenose dolphins (*Tursiops truncatus*) housed at Dolphin Quest Oahu (Honolulu, HI, USA) with an average (± S.D.) age of 22.8±9.9 years (range = 11 – 35 years) and body mass of 198.1±42.9 kg (range = 147.0 – 251.7 kg, **Table 1**) participated in this study. All data were collected under voluntary participation and the animals could end a trial at any time. Routine veterinary assessments include venous blood sampling, and the dolphins that participated in this study had previously been desensitized to the blood sampling protocol. The study protocols were accepted by Dolphin Quest and the Animal Care and Welfare Committee at the Oceanogràfic (OCE-17-16, amendments OCE-29-18 and OCE-3-19i).

**Table 1.**
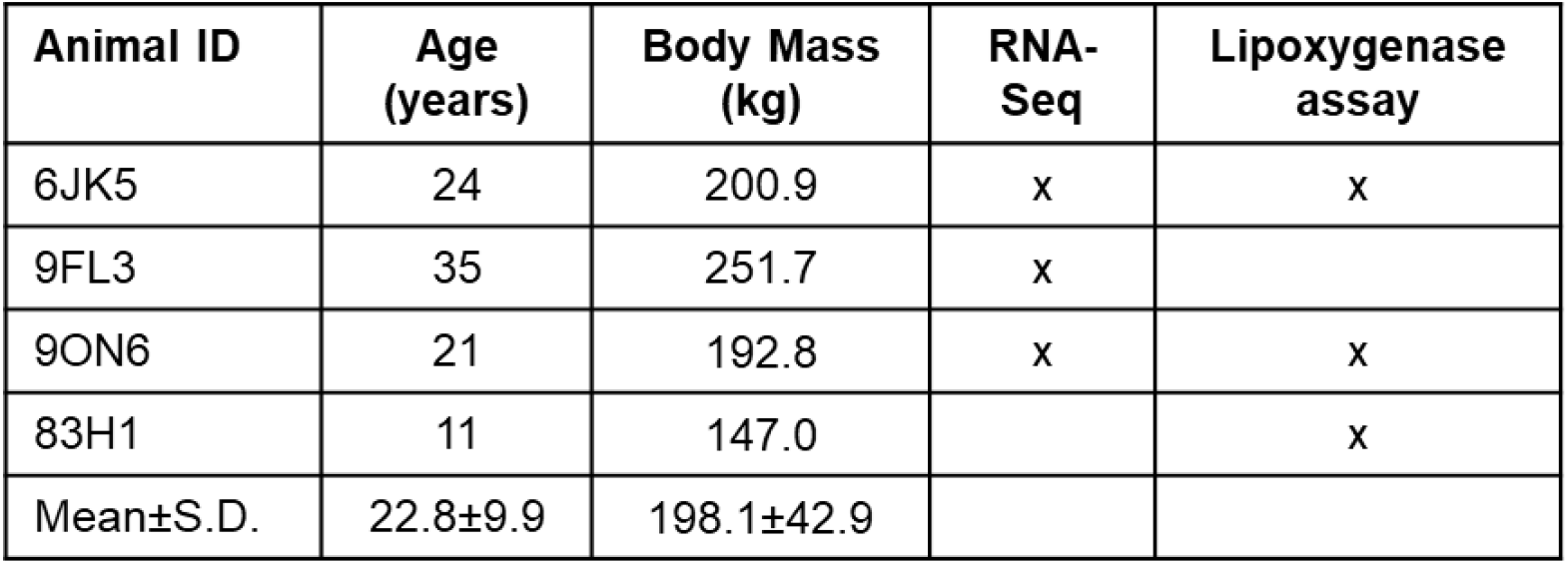
Animal ID, age (years), body mass (kg), and included analyses for all dolphins in the study.

### Experimental trials

Serum was isolated from whole blood samples at baseline, 3 minutes, and 4 ½ − 5 minutes of breath-holding on fasted dolphins at Dolphin Quest, Oahu, March 2018 and May 2019. All trials were performed in the morning, when the animals were fasted with at least 12 hours having passed since the last meal on the previous day to minimize the potential confounding effect of nutritional state. To ensure that the samples were collected during resting behavior each breath-hold was proceeded by 2 minutes of rest or slow swimming at the surface. A trial was initiated when the dolphin rolled into dorsal recumbency with its blowhole submerged and continued for approximately 5 minutes. The breath-hold ended when the animal rolled into ventral recumbency and took a breath. Prior to this study the animals had previously participated in breath-hold experiments of durations up to 5 minutes (Fahlman et al., 2019a, 2020b).

### Blood collection and processing

Whole blood was collected from tail flukes at baseline (0-30 seconds into the breath-hold) and during breath-holding for 3 minutes and 4 ½ (2018) or 5 (2019) minutes while the animal was in dorsal recumbency with its blowhole submerged (**Figure 2A**). Blood was collected into PAXgene tubes and RNA-Seq was performed subsequent to shipping, red blood cell lysis, RNA extraction (**Figure 2A**). All samples were shipped the same day via overnight courier to Duke University for downstream processing. For RNA extraction, tubes were equilibrated to room temperature for 2 hours to achieve complete lysis of blood cells. Subsequently, tubes were centrifuged at 4,000 x g for 10 minutes. Pellets were resuspended in 4 mL of RNase-free water and RNA was extracted according to the PAXgene Blood RNA kit (PreAnalytiX #762164). Prior to library prep, RNA quality was evaluated on a Bioanalyzer 2100 (Agilent). Stranded mRNA-seq libraries were prepared using the Nugen Universal Plus mRNA-seq Library preparation kit with Globin AnyDeplete (Tecan #9147-A01). Libraires were sequenced at 150bp paired-end on one lane of an Illumina NovaSeq 6000 instrument S-Prime flow cell. Library preparation and sequencing was performed in conjunction with the Duke University Sequencing and Genomic Technologies Shared Resource. Samples collected in 2018 were used to conduct RNA-Seq analysis and samples collected in 2019 were used for the lipoxygenase assays.

**Figure 2.**
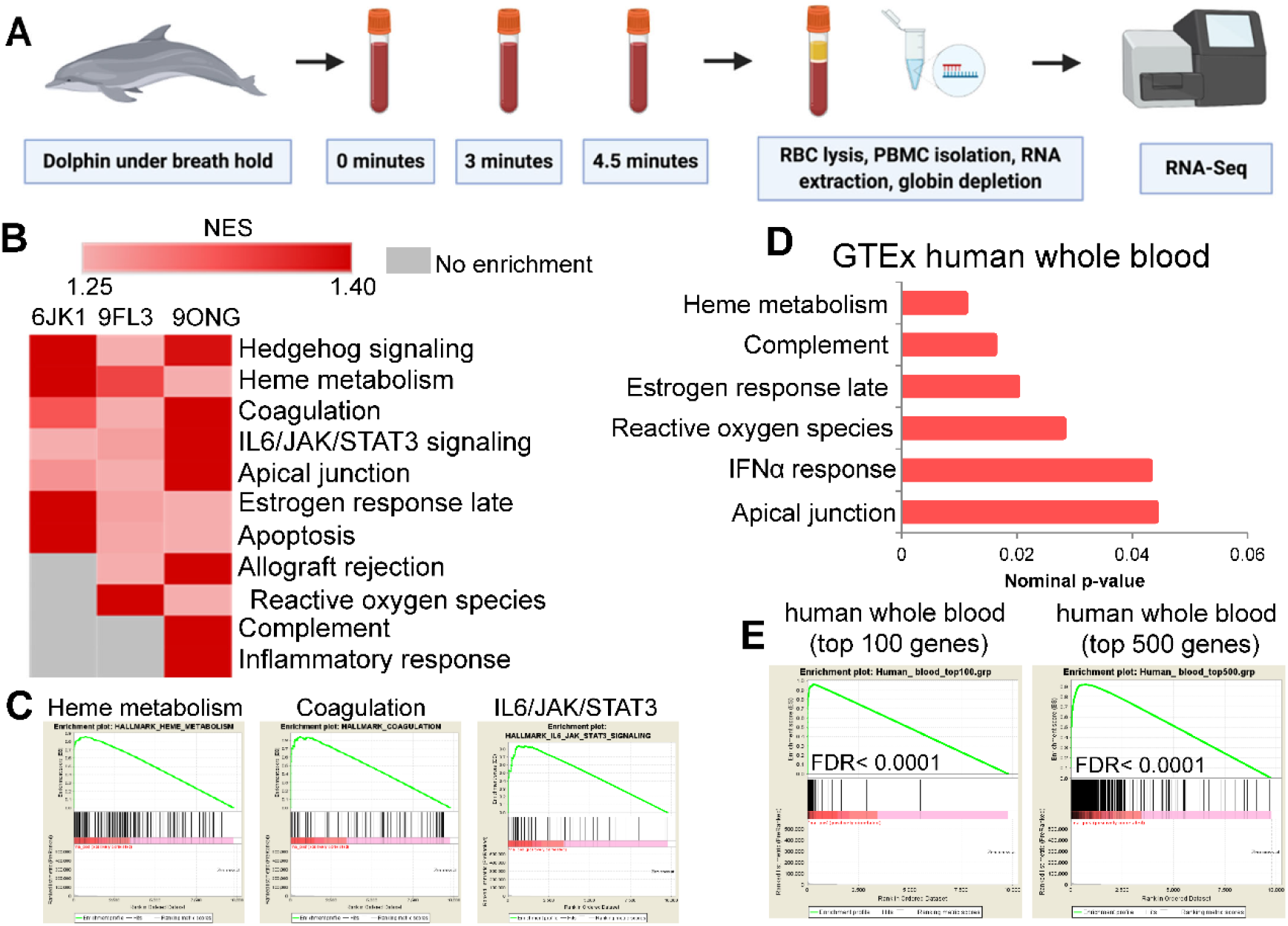
RNA-Seq from dolphin peripheral blood mononuclear cells reveals enrichment of pathways similar to humans. A. Whole blood from dolphins undergoing fasted breath-holds at baseline (0-30 seconds), 3 minutes, and 4 ½ minutes was collected from tail flukes and stored in PAXgene tubes for RNA extraction of peripheral blood mononuclear cells and RNA-Seq. B. Gene set enrichment analysis of baseline RNA-Seq data ranked by total expression pinpoints highly expressed relevant pathways. C. Enrichment plots for heme metabolism, coagulation, and IL6/JAK/STAT3 signaling from baseline dolphin RNA-Seq data. D. GSEA-based pathway enrichment from GTEx human whole blood RNA-Seq data ranked by total expression. E. GSEA enrichment plots comparing dolphin RNA-Seq data ranked by total expression with top 100 and top 500 expressed genes in human whole blood.

### RNA-Seq data analysis

RNA-seq data was processed using the TrimGalore toolkit (Krueger, 2020) which employs Cutadapt (Martin, 2011) to trim low-quality bases and Illumina sequencing adapters from the 3’ end of the reads. Only reads that were 20nt or longer after trimming were kept for further analysis. Reads were mapped to the turTru1v92 version of the dolphin genome and transcriptome (Kersey et al., 2012) using the STAR RNA-seq alignment tool (Dobin et al., 2013). Reads were kept for subsequent analysis if they mapped to a single genomic location. Gene counts were compiled using the HTSeq tool (Anders et al., 2015). Only genes that had at least 10 reads in any given library were used in subsequent analysis. Normalization and differential expression across the time points were carried out using the DESeq2 (Love et al., 2014) Bioconductor (Huber et al., 2015) package with the R statistical programming environment (R Core Team, 2020). The false discovery rate was calculated to control for multiple hypothesis testing. To identify relevant molecular features of dolphin breath-holding we first analyzed the RNA-Seq data from all individuals at baseline using gene set enrichment analysis (GSEA) (Mootha et al., 2003; Subramanian et al., 2005). GSEA is a standard pathway analysis tool that calculates enrichment scores for annotated pathways based on the rank order of genes present in the data for each pathway. Pathways with genes that are more up- or down-regulated are more likely to be enriched in a data set than pathways whose genes are randomly distributed throughout the data. Pathway enrichments in dolphin PBMCs at baseline, with genes ranked on total expression value, were compared with human whole blood pathway enrichments from the Genotype-Tissue Expression (GTEx) project.

### Construction of gene regulatory networks

Gene expression networks were created using GeneMANIA (Franz et al., 2018), implemented within the Cytoscape platform (Shannon et al., 2003). For time-dependent gene network construction all nodes with 0 or 1 connection were trimmed out of the networks. Two additional non-coding RNA genes were eliminated (RF00016 and RF00026). To quantify network connectivity, all genes in the network were individually ranked by the following network parameters: degree, clustering coefficient, closeness, betweenness, neighborhood connectivity, and stress. These rankings were summed to generate a sum rank score for each gene. Pathway enrichments were performed in STRING using the trimmed network of 123 genes. Human whole blood transcriptomics data used for the analyses described in this manuscript were obtained from the Genotype-Tissue Expression (GTEx) Program Portal (https://gtexportal.org/home/) accessed on 9/20/2020.

### Lipoxygenase assays

Briefly, 5 ml of blood was collected directly into BD Vacutainer® SST™ Tubes (SST) using a 21 g, ¾ in. winged infusion set with a BD Vacutainer adapter and holder. Tubes were gently inverted 5 times to activate clotting reagent and allowed to clot at room temperature for 30 minutes in an upright position. Tubes were centrifuged at 1,500 x g for 15 minutes to separate serum fractions, and serum was transferred to 15 ml conical tubes, frozen on dry ice, and shipped to Duke University for downstream analyses. Sera were stored at −80°C until use. Lipoxygenase activity was quantified from 1 μg of total protein using a Fluorometric Lipoxygenase Activity Assay Kit (BioVision Inc; cat. #K978).

## Results

### RNA-Seq from dolphins at baseline pinpoints enriched gene regulatory networks

All samples produced between 30 and 40 million reads, with no time-dependent changes in read counts across samples (**Supplementary Figure 1A**). Principle components analysis and hierarchical clustering of all samples (three individual dolphins x three time points) revealed both individual- and within-individual time-dependent grouping of the data (**Supplementary Figure 1B, C**). GSEA identified multiple pathways enriched in dolphin PBMCs at baseline when ranked by total expression, including hedgehog signaling and several pathways relevant to blood cell metabolism, including heme metabolism, coagulation, IL6/JAK/STAT3 activation, apical junctions, and allograft rejection (**Figure 2B, C**). GSEA also identified enrichment of pathways related to apical junctions, interferon alpha response, estrogen response, complement activity, and heme metabolism in RNA-Seq data from GTEx human whole blood transcriptomes (**Figure 2D**). Comparison of dolphin baseline RNA-Seq data ranked by total expression with the top 100 and 500 most highly-expressed genes in human whole blood showed significant enrichment (FDR<0.0001) (**Figure 2E**). Together these analyses suggest that significant overlap exists in mRNA expression at both the gene-level and pathway-level between dolphin and human blood.

### Breath-holding induces upregulation of multiple regulatory pathways

We next reasoned that patterns of step-wise increases in mRNA expression may pinpoint molecular responses to breath-holding common across individuals. We constructed gene regulatory networks for 136 genes with step-wise increases in mRNA expression from baseline to 3 minutes and again from 3 minutes to 4 ½ minutes (**Figure 3A**). We performed network analysis to identify genes that are upregulated and have the most network interactions. To do this we analyzed the time-dependent gene regulatory network for the following parameters: degree, clustering coefficient, closeness, betweenness, neighborhood connectivity, and stress. We then plotted the sum rank score of these network parameters with gene expression for each gene in the network. These analyses pinpointed arachidonate 5-lipoxygenase (ALOX5) as among the most connected genes with a time-dependent increase in expression (**Figure 3B, C**). Additional genes, including EPX, PTGDR2, SIX5, DCN, ADAMTS12, and GLRX2 demonstrated upregulation and/or high network connectivity (**Figure 3B, C**). We used GeneMANIA to infer transcription factor and microRNA targets from this time-dependent network. The gene regulatory network produced from these genes displayed enrichment in targets from several transcription factor families, including GATA and the <u>s</u>mall, <u>m</u>others <u>a</u>gainst <u>d</u>ecapentaplegic (SMAD) families (**Figure 3D**), both of which have been implicated in hematopoietic development and regulation (Blank and Karlsson, 2011; Lentjes et al., 2016). Network inference also pinpointed enrichment of targets of multiple microRNAs, including the miR148A/B/152 family, miR492, miR186, miR518A-2, the miR130A/B/301 family, and miR205 (**Figure 3E**). Consistent with the identification of ALOX5 as a core network node, the network was functionally enriched the synthesis of 5-eicosatetraenoic acid pathway, which is an initial step in the production of arachidonic acid by ALOX5 (**Figure 3F**).

**Figure 3.**
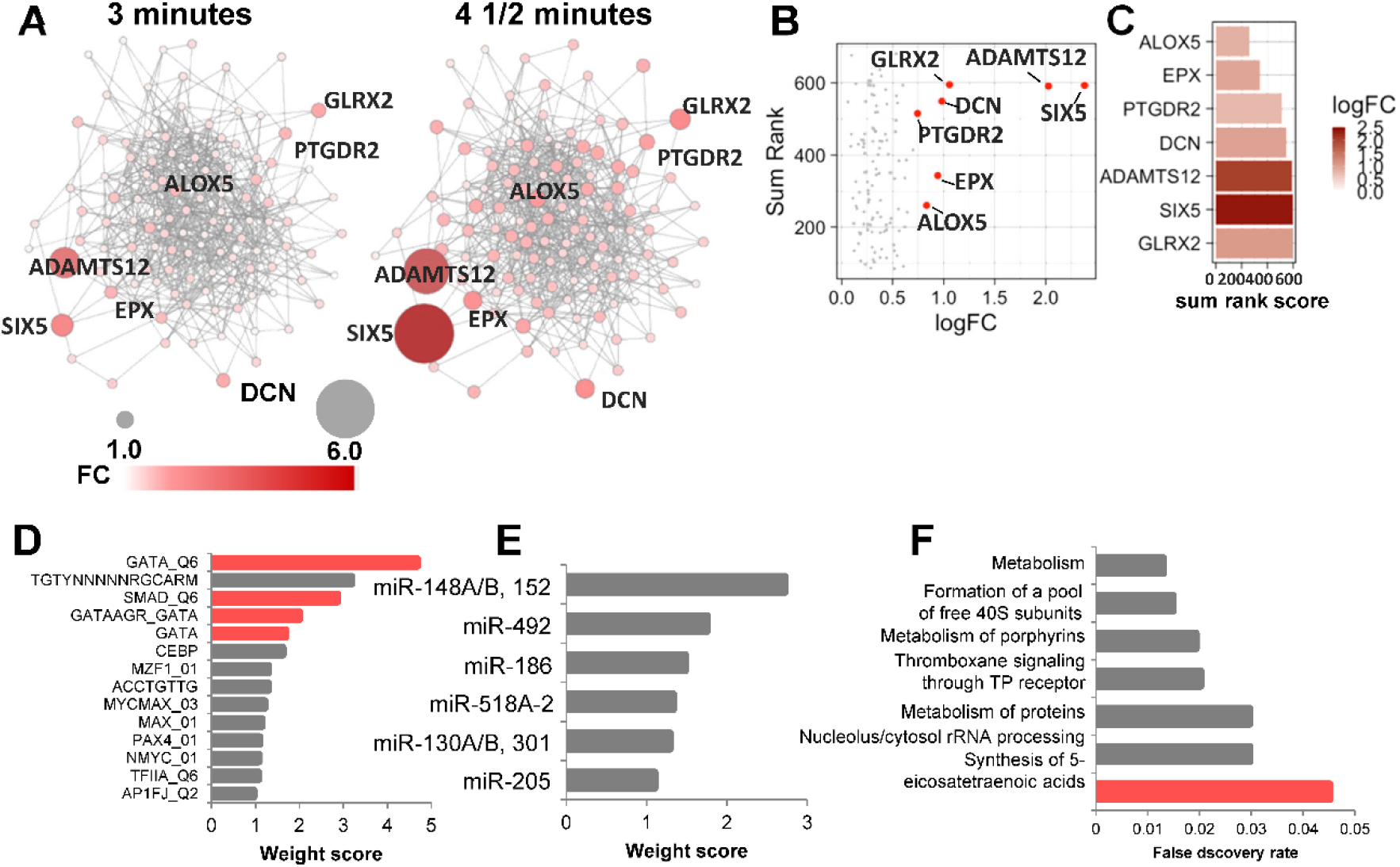
Time-dependent upregulation of gene regulatory pathways during dolphin breath holding. A. Gene regulatory network formed by the time-dependent increases in mRNAs from baseline to 3 minutes and 4 ½ minutes. Fold changes for each gene over time are indicated by darker red and larger nodes. B. Network analysis of genes within a co-expression network with increased expression over time. C. Top genes with increased expression sorted by their network analysis parameters. D. GeneMANIA-based transcription factor inference pinpoints GATA and SMAD transcription factor targets within the time-dependent network. E. MicroRNA enrichment inference based on the time-dependent network. F. Functional pathway enrichments for the time-dependent gene regulatory network.

### Arachidonate 5-Lipoxygenase (ALOX5) and subsequent lipoxygenase activity enhanced in breath-holding dolphins

Based on the network analyses we focused on ALOX5 for further validation. At the mRNA level, ALOX5 was one of just two genes, along with IL5RA, that was significantly upregulated in all three individuals during breath-holding (**Figure 4A, B**). Lipoxygenase assays from serum of three individual dolphins collected in 2019 revealed time-dependent increases in lipoxygenase activity during breath-holding in all three individuals, consistent with the RNA-Seq analyses (**Figure 4C**). Comparison of the timing of these molecular changes with previously-published *f*_H_ measurements from the same dolphins demonstrated that changes in gene expression and enzymatic activity were coincident with the expected timing of bradycardia based on the heart rate data (**Figure 4D**). Overlay of the RNA-Seq data for ALOX5 expression with the heart rate data shows the upregulation of ALOX5 is concomitant with lower heart rate (**Figure 4E**).

**Figure 4.**
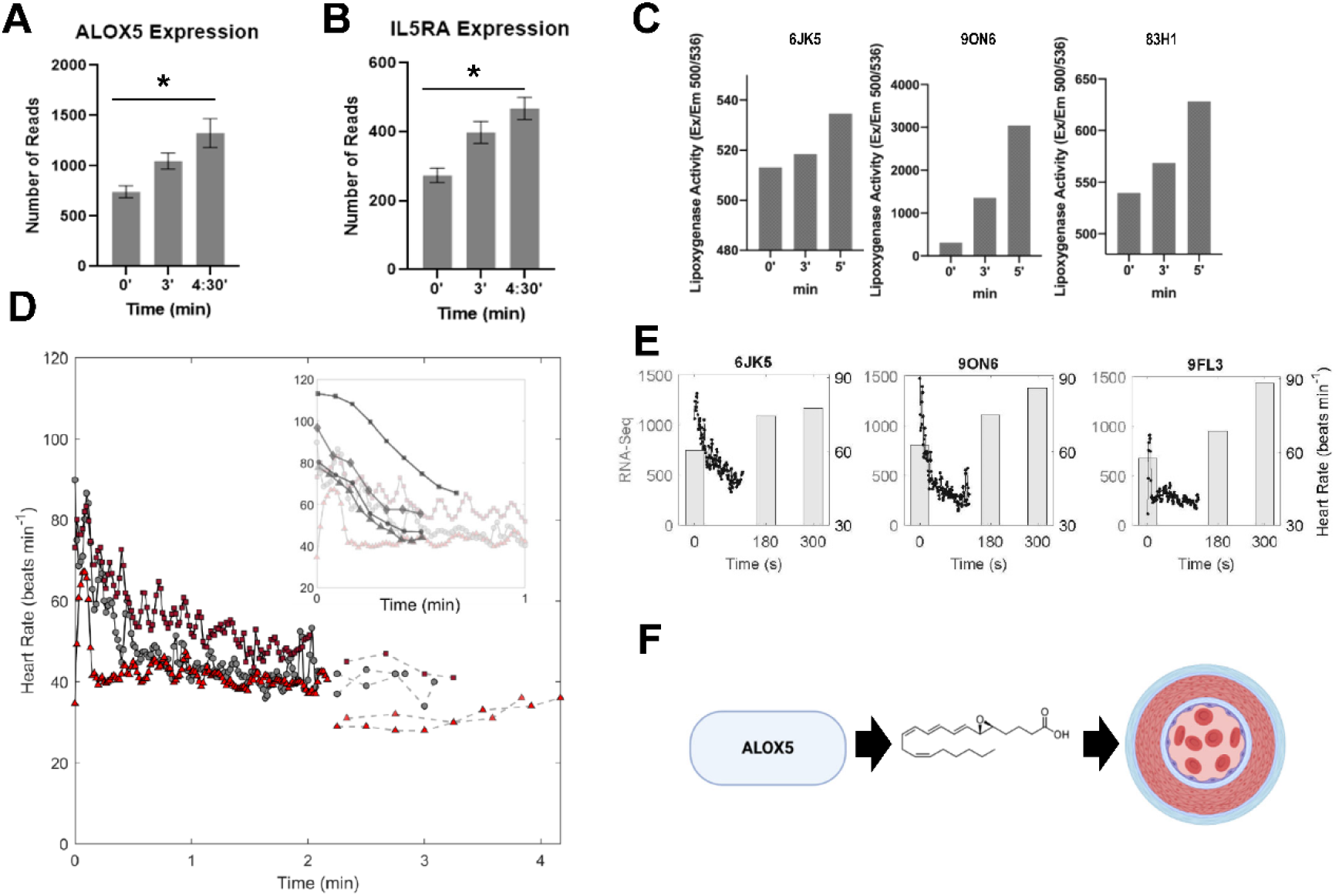
Dolphins induce ALOX5 activity during breath-holding. A. ALOX5 and B. IL5RA mRNA expression is significantly increased over time during breath-holding. C. Individual dolphin lipoxygenase activity in whole blood collected at an independent sampling date. D. Physiological measurements of heart rate for three individual dolphins (black lines from ECG data previously published in Blawas et al., 2020, dashed lines from echocardiogram data previously published in Fahlman et al., 2020) over time. Inset shows heart rate for humans performing breath-holds with facial immersion in water (dark gray in inset) overlaid on dolphin heart rate. Human heart rate traces were digitally extracted from Arnold, 1985; Andersson et al., 2004; and Shattock and Tipton, 2012. E. Overlay of heart rate data with ALOX activity in three individual dolphins. F. Hypothesized mechanism through which ALOX5 improves dive performance and mitigates ischemic stress tolerance.

## Discussion

Dolphins and other cetaceans have evolved exquisite physiological adaptations to deal with the challenges of a fully aquatic lifestyle including having a hydrodynamic shape to reduce drag (Fish, 1993), counter-current heat exchangers for thermoregulation (Favilla and Costa, 2020; Pabst et al., 1999), and cardiorespiratory plasticity for exquisite management of circulation and respiratory gases (Blawas et al., 2021; Fahlman et al., 2020b, 2020a, 2019b; Noren et al., 2012). The well-known dive response, a suite of adaptations that support reduced aerobic metabolism during diving, involves apnea, bradycardia, and peripheral vasoconstriction that assures maintained mean arterial blood pressure as blood flow to peripheral tissues is reduced and allows regulation of perfusion to conserve oxygen-rich blood for the brain and heart. To maintain a constant mean arterial blood pressure and prevent hypertension, these adaptations must work in concert to ensure efficient autoregulation; however, extended dives also result in frequent events of ischemia and hypoxia (Fahlman et al., 2019a; McKnight et al., 2019; Ridgway et al., 1969). While these cardiorespiratory adaptations have been studied from the perspective of the changes in *f*_H_ associated with diving, we are not aware of any study that has measured blood pressure in voluntarily diving cetaceans. Thus, little is known about whether dolphins are able to maintain constant mean arterial blood pressure throughout the breath-hold. In addition, knowledge of the molecular adaptations that contribute to enhanced tolerance to hypoxia and ischemic stress, and that prevent reperfusion injury during and following a dive, is rudimentary at best. To address this lack of understanding, we combined an integrated genomics and systems-level analysis of breath-hold responses at the molecular level with existing physiological measurements to define the molecular responses to breath-holding in dolphins.

While this study is limited by a small sample size and relatively short breath-hold durations, our analyses identified candidate genes and pathways with time-dependent changes in expression throughout the breath-holds that were validated in functional studies using independently-collected samples and assays. Consequently, these results provide evidence for fine-scale cardiovascular control in bottlenose dolphins at the genetic level, and suggest that dolphins may manage blood pressure changes during diving using both autonomic and molecular pathways to regulate peripheral vasomotor control. Notably, these molecular changes occurred within the calculated aerobic dive limit (cADL) of bottlenose dolphins - the duration of a dive that can be sustained without requiring anaerobic respiration at the cellular level which has been estimated to be 6.5 minutes (Fahlman et al., 2018). This suggests that changes in gene expression may operate on a short enough time-scale that they could play a role in driving the physiological changes that are observed during the breath-hold and recovery. It is also worth considering the possibility that changes in gene expression could occur to support specific physiological responses to diving during a dive, and that this gene expression differs when the animal is at the surface. Future studies will be focused on using novel technologies, such as GRO-Seq (Lopes et al., 2017) and others to measure nascent mRNAs, as well as measuring later time points to understand the changes that occur upon recovery from breath-holds.

To provide physiological context for these molecular alterations on the time scales observed, we compared molecular changes to changes in previously published *f*_H_ patterns in the same individual dolphins during submerged breath-holds (Blawas et al., 2021; Fahlman et al., 2020b). If we assume that the appearance of vasoconstriction is coincident with bradycardia, our data provide evidence of an increase in the expression of a gene, ALOX5, known to promote vasoconstriction coincident with the onset of vasoconstriction. Vasoconstriction, or a narrowing of the blood vessels, has been suggested as a mechanism by which marine mammals during forced dives have been observed to optimize the use of onboard oxygen stores in the blood and muscle (Davis and Kanatous, 1999; Scholander and Grinnell, 1942; Zapol et al., 1979). Given the long assumed link between vasoconstriction and bradycardia in marine mammals, the rapid bradycardia we observed suggests that vasoconstriction was occurring in the dolphins in this study during breath-holds (Hochachka, 1981; Van Citters et al., 1965). We found that changes in gene expression occurred in all animals during the 5-minute breath-hold trials and that the same gene families that were upregulated in the dolphins during breath-holds help manage vasoconstriction in mice (Ichinose et al., 2001) and humans (Friedman et al., 1984).

Our integrated approach reveals possible molecular underpinnings that may support and act synergistically with the cardiac response to breath-holding in bottlenose dolphins. Specifically, we identified a suite of candidate genes that may support peripheral vasoconstriction and provide defense against ischemic and hypoxic stress in dolphins, including the GATA and SMAD transcription factors, several microRNAs, a disintegrin and metalloproteinase with thrombospondin motifs 12 (ADAMTS12), mitochondrial glutaredoxin-2 (Glrx2), and ALOX5. Interestingly, many of these factors play known roles in regulating hypoxia, hematopoiesis, and ischemic stress responses. For example, the GATA transcription factor family is an important modulator of hematopoietic development of T lymphocytes, mast cells, and erythrocytes (Lentjes et al., 2016). Likewise, the SMAD family regulates hematopoietic stem cells (Blank and Karlsson, 2011). Of the microRNAs identified from our analysis of target enrichments, nearly all have been shown to be protective against ischemia-induced cell death, including miR148A (Zheng et al., 2018), miR492 (Guo et al., 2020), miR186 (Bostjancic et al., 2009; Li et al., 2013; Wang et al., 2018), miR130 (Lu et al., 2015), and miR205 (Chen et al., 2019). At the protein-coding gene level, ADAMTS12 genetic variation is associated with pediatric stroke (Witten et al., 2020), GLRX2 is implicated in neuroprotection during hypoxia and ischemia (Romero et al., 2015), and ALOX5 is known to be induced by hypoxia (Porter et al., 2014) and mediates the production of leukotrienes, which induce bronchoconstriction and vasoconstriction (Poeckel and Funk, 2010). In addition, both ALOX5 and IL5RA have been identified as susceptibility genes associated with asthma and asthmatic inflammation in humans (Cheong et al., 2005; Mougey et al., 2013), and a monoclonal antibody to the IL5RA ligand, IL5, is FDA-approved for the treatment of severe eosinophilic asthma (Fala, 2016; Pavord et al., 2012). Given the intricate connection between molecular control and physiologic function to manage ischemia, hypoxia, and inflammatory responses in humans and rodent models, (Bartels et al., 2013) it is intriguing to speculate as to how dolphins and other marine mammals may uncouple or leverage these interconnected processes for improved tolerance to ischemic/hypoxic stress without the pathological consequences associated with hyper-stimulation of these processes.

These results demonstrate that the ALOX5 pathway is upregulated in bottlenose dolphins during breath-holds and offer a potential mechanism for maintaining elevated peripheral resistance through vasoconstriction, which helps manage blood distribution and the available oxygen for critical organs. We suggest that the upregulation of ALOX5 could support a genetic response that is secondary to the autonomic response during diving to prolong vasoconstriction and maintain mean arterial blood pressure during extended periods of submersion (**Figure 4F**). Interestingly, the changes we observed occurred within the cADL of the species, indicating that fluctuations in gene expression could be occurring during regular dives. These fast-acting changes in gene expression that support vasoconstriction provide evidence for fine-scale control of perfusion in dolphins and an ability to maintain constant blood pressure, as is observed during forced dives of pinnipeds (Blix et al., 1976; Zapol et al., 1979). Additionally, the data show that during the breath-holds a large suite of candidate genes are upregulated that may support an increased tolerance to the hypoxia and ischemic conditions that are expected to arise in some peripheral tissues during diving. By interpreting these molecular data in the context of the physiological changes known to occur in bottlenose dolphins during a breath-hold, we have identified several genes that are upregulated during apnea in dolphins and may be an additional mechanism to reinforce vasoconstriction while also providing defense against the hypoxia and ischemia resulting from this response.

By examining molecular data through a physiological lens, these data connect the cellular and tissue-level responses of dolphins to apnea to understand whether the bottlenose dolphin may be genetically tuned to dive and withstand the hypoxia and the potential implications of this to translational medicine. Our results uncover potential candidates at the intersection of ischemia, hypoxia, and vasoconstriction that may contribute to the exquisite adaptation of dolphins and other marine mammals to life in the ocean.

## Abbreviations List

cADL: calculated aerobic dive limit
ALOX5: Arachidonate 5-Lipoxygenase
GSEA: Gene Set Enrichment Analysis
*f*_H_: heart rate
IL5RA: Interleukin 5 receptor, alpha
PBMC: peripheral blood mononuclear cells

## Acknowledgments

The authors wish to thank the marine mammal specialists, veterinarians, and dolphins at Dolphin Quest, Oahu, The Duke Sequencing and Genomics Technologies Shared Resource, and the Duke Genomics Analysis and Bioinformatics Shared Resource. The authors also thank the Laboratory of Dr. Peter Hoffman at The University of Hawai’i for providing a centrifuge and other equipment for sample processing. This work was supported through Dolphin Quest, Oahu (JAS) and the Triangle Center for Evolutionary Medicine (JAS). AMB was supported by a Bureau of Ocean Energy Management (BOEM) Environmental Study and the E. Bayard Halsted Scholarship in Science, History, and Journalism. The authors would like to thank Giselle Vargas and Mallissa Vuong for their role in data collection.

