## Supplemental Figure 1 for "An integrated comparative physiology and molecular approach pinpoints mediators of breath-hold capacity in dolphins"

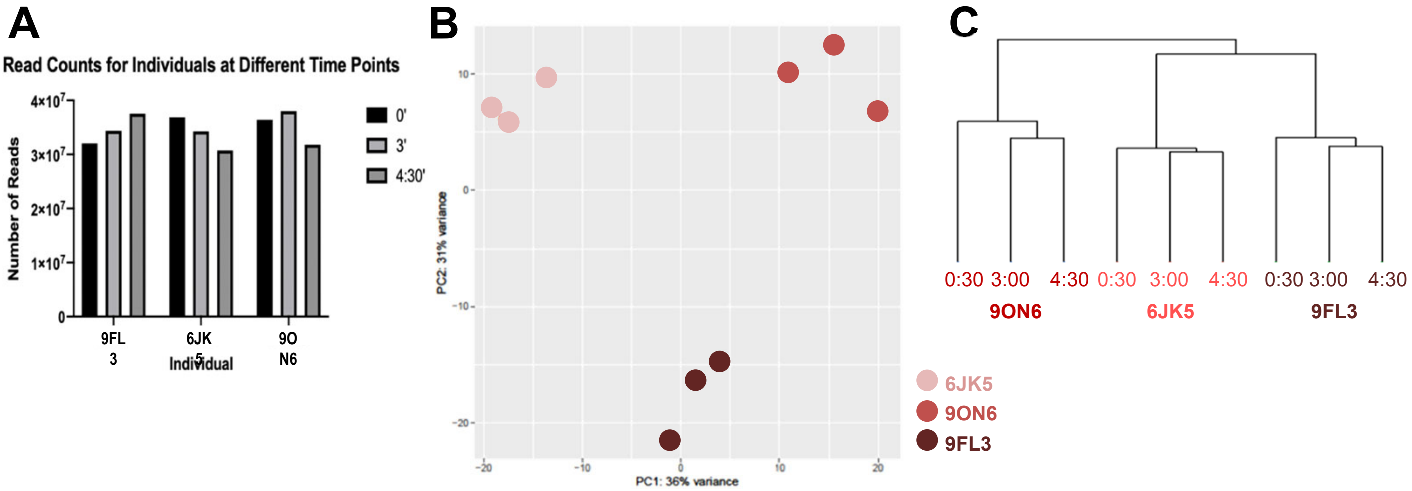


**Fig. S1.** RNA-Seq on peripheral blood mononuclear cells from dolphin breath holding suggests individual and time-dependent signals. **A.** Total read counts for all individuals and time points. **B.** Principal components analysis shows clustering of samples by individual dolphin. **C.** Hierarchical clustering reveals time-dependent signals for each individual, with the baseline sample from each individual the most basal and the two later time points clustered together.
